# Degradation and release of dissolved environmental RNAs from zebrafish cells

**DOI:** 10.1101/2023.07.25.550455

**Authors:** Zhongneng Xu

## Abstract

Environmental RNAs in water are gradually being applied in aquatic ecological surveys, water pollution monitoring, etc., but the current methods to detect environmental RNAs in water can mainly measure the RNAs in the filters that are used for filtering water samples, neglecting dissolved environmental RNAs in water. The sources and degradation profiles of dissolved environmental RNAs in water remain unknown. The present study was conducted to measure the permeability of extracted RNAs from zebrafish cells through filters, the degradation of extracted RNAs from zebrafish cells in tubes, and the release rate and degradation of dissolved environmental RNAs from living zebrafish cells and dying zebrafish cells, aiming to provide dynamic information from dissolved environmental RNAs in water. The results showed that there were no significant differences between the levels of extracted RNAs from zebrafish cells before filtration with 0.45 µm filters and those in the filtrates. The extracted RNAs from zebrafish cells degraded in water in the tubes, and after 2 months, more than 15% of RNAs in the groups of RNAs in water were still detected. The half-life of all the RNAs in the tubes was approximately 20∼43 days. During the 6-day experiment of the release and degradation of dissolved RNAs from living cells, an average of 4.1×10^-4^ ∼ 1.7×10^-3^ pg dissolved RNAs (7.6×10^5^ ∼ 3.2×10^6^ RNA bases) were secreted per cell per day into the liquid environment. During the 6-day experiment of the release and degradation of dissolved RNAs from dying cells, approximately 4.2 pg of dissolved RNAs released by a dying zebrafish cell in water could be detected. The dissolved environmental RNAs in water from zebrafish cells degraded faster in the presence of zebrafish cells: under the conditions without zebrafish cells, the average survival rate of the dissolved environmental RNAs in water per day was 98.4%/day; under the conditions with living zebrafish cells, the average survival rate per day was 49.7%/day; and under the conditions with dying zebrafish cells, the average survival rate per day was 34.9%/day. The estimated levels of dissolved environmental RNAs in water in fish tanks were too low to be detected by the current techniques. Although the methods in the present study need to be improved, this study may provide information to develop new ways to measure the dynamics of dissolved environmental RNAs in water and quantitatively analyze RNAs released into liquid environments of living and dying cells.

## 1. Introduction

Environmental RNAs are an exciting and expected evaluation indicator in ecological and environmental research fields (Laroche et al., 2017; Pochon et al., 2017; Cristescu, 2019; Veilleux et al., 2021; Yates et al., 2021). The detection of environmental RNAs in water is increasingly applied in fishery resource surveys, aquatic ecological biodiversity surveys, water pollution monitoring, and the analysis of the living state of aquatic species (Foley et al., 2016; Pochon et al., 2017; Wood et al., 2020; Marshall et al., 2021; Miyata et al., 2021; Yates et al., 2021; Jo et al., 2022b; Kagzi et al., 2022; Littlefair et al., 2022; Robert et al., 2022).

However, the current filter methods to detect environmental RNAs in water can mainly measure RNAs remaining in the filters, neglecting dissolved RNAs in the filtrates. Filters used for filtering water samples to measure environmental RNAs have different pore sizes, such as 0.2∼0.22 µm (Wu and Liu, 2018; Wood et al., 2020; Hempel et al., 2022), 0.45 µm (von Ammon et al., 2019; Tsuri et al., 2020; Miyata et al., 2021; Zaiko et al., 2022; Jo et al., 2022b; Miyata et al., 2022), 0.7 µm (Kagzi et al., 2022; Hechler et al., 2022; Jo et al., 2022a; Littlefair et al., 2022), 1.2 µm (Marshall et al., 2021; Zaiko et al., 2022), 1.6 µm (Pochon et al., 2017), and 5 µm (Zaiko et al., 2022). A study indicated that filters with different pore sizes could collect naked RNAs (Zaiko et al., 2022), but the RNA amounts in the filtrates were unknown in that study. Dissolved RNAs can pass through the filters, and the RNA amounts in the filtrates can be detected (Sakano and Kamatani, 1992). Although some parts of dissolved RNAs of water samples might attach to the filters or mix with the RNAs in suspended particles in the filters, the actual amounts and fluctuation trends of dissolved environmental RNAs in water and the sources of dissolved environmental RNAs in water are still unclear.

Therefore, the present study conducted experiments to examine the level fluctuations of dissolved RNAs in water released by zebrafish cells under laboratory conditions, with the aim of exploring some dynamic information and sources of dissolved environmental RNAs in water.

## 2. Results

### 2.1 Free RNAs through the 0.45 µm filter

The results showed that 87% of RNAs remained in the filtrates (Figure 1), and there were no significant differences between RNA levels in the samples before and after filtration (P>0.05). There were no significant differences between the levels of RNAs with different sizes, such as 18S RNAs and 28S RNAs, in the samples before and after filtration (P>0.05). That is, from a statistical point of view, all sizes of RNAs exacted from zebrafish cells could pass through the 0.45 µm filters.

**Figure 1.**
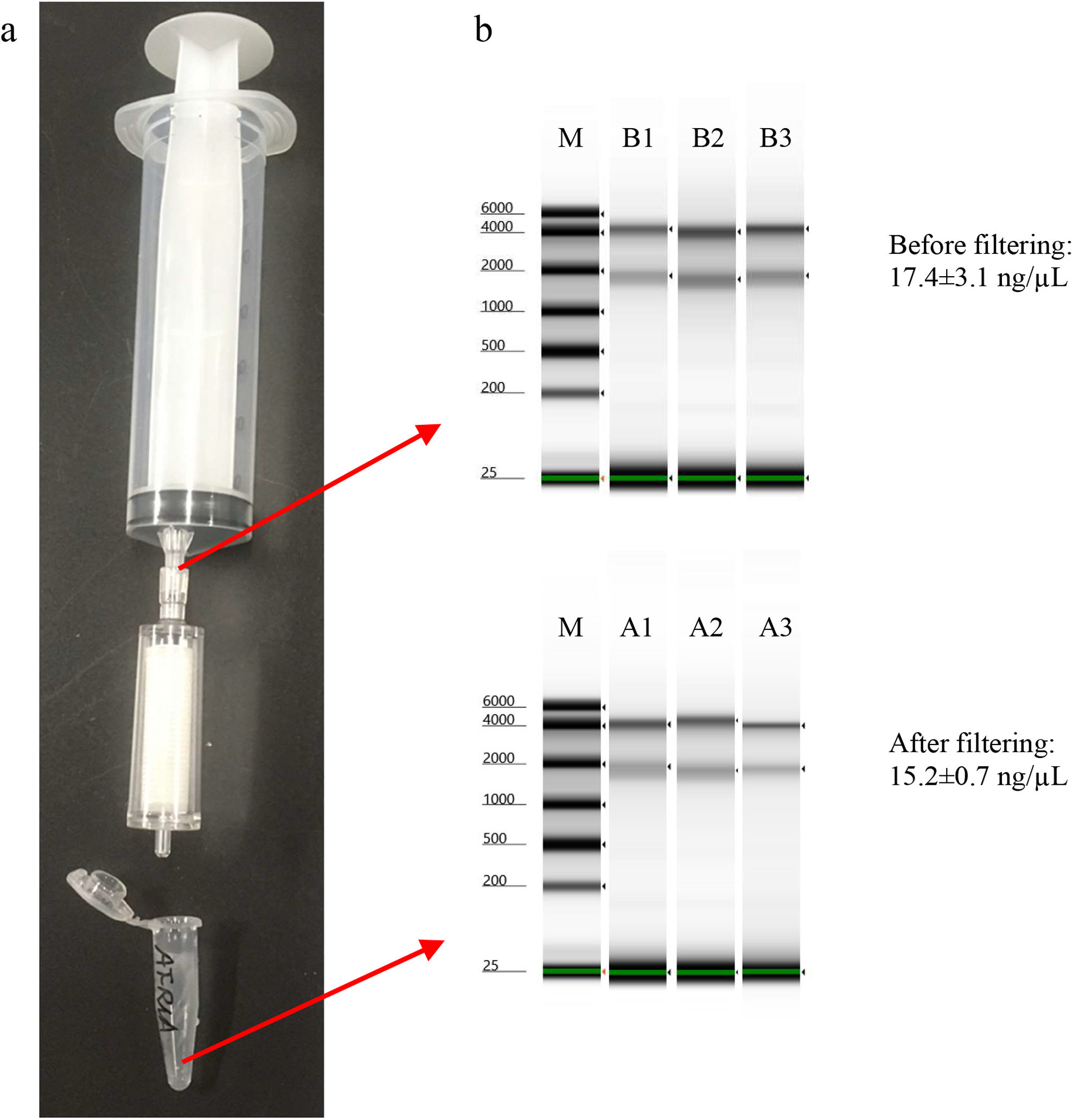
Free RNAs in water through the 0.45 µm filter. a. The device for filtering liquid samples and collecting the filtrates. The pore size of the filter unit was 0.45 µm. b. The gel results of RNA ScreenTape. M represents the marker. B1, B2, and B3 are the repeats of original samples, in which the extracted RNAs from zebrafish BRF41 cells were added before filtering. A1, A2, and A3 are the samples after filtering with 0.45 µm filter units.

### 2.2 Degradation process of free RNAs in water in the tubes

The degradation process of RNAs extracted from zebrafish cells in an aqueous environment in centrifuge tubes could be measured by the TapeStation system, gel electrophoresis, and NanoDrop method (Figure 2). After 2 months, more than 15% of RNAs in the groups of RNAs in water were still detected, while the RNA levels of the water control groups were very low (3.2∼3.7 ng RNA/μL) during the two-month experimental period. Over time, RNAs with long sequences were broken down into short and middle RNA fragments (Figure 2a, Figure 2b). After two months, detectable RNAs in the samples were almost all short fragments. These results might indicate that the short RNAs in water had long lifespans. The half-life of RNAs longer than 25 bases in the tubes detected by the TapeStation system was approximately 20∼30 days (Figure 2a), and the half-lives of all the RNAs in the tubes detected by NanoDrop were approximately 35∼43 days (Figure 2c).

**Figure 2.**
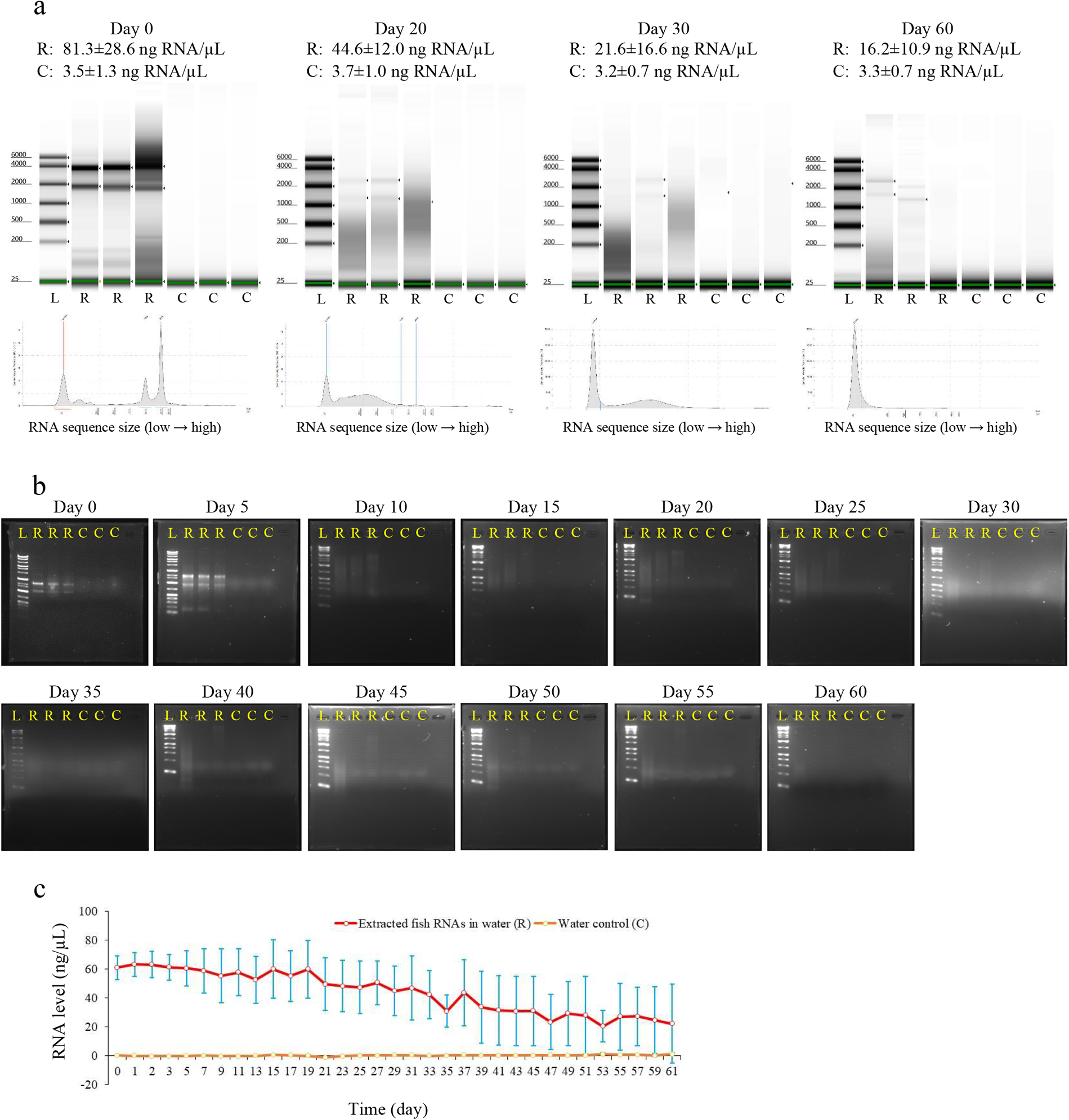
Degradation process of dissolved RNAs in water in the centrifuge tubes. In the figure, L represents the marker or the ladder, R represents extracted zebrafish RNAs in water, and C represents the water control. a. The results of detection by the TapeStation system. b. The results of the detection by normal gel electrophoresis. c. The results of the detection by NanoDrop. The solid red curve represents extracted zebrafish RNAs in water, the dashed orange curve represents the water control, and the blue bar is the standard deviation. The RNA level of the water control is so low that the curve of the water control overlaps the abscissa.

### 2.3 Dissolved environmental RNAs released by living cells

The rates and total amount of dissolved RNAs released from the living zebrafish cells in L-15 medium were detectable by the present method (Figure 3). During the experiment, an average of 0.41∼1.7 pg dissolved RNAs (approximately 1.7∼7.2% cellular RNAs) were secreted per cell per day into the liquid environment. If the single nucleotide in RNA is 325 g/mole on average, during the experiment, an average of 7.6×10^8^ ∼ 3.2×10^9^ RNA bases was secreted per cell per day into the liquid environment; thus, there may be rich biological information in these bases. The sizes of the dissolved RNAs released covered a large scale, from less than 200 bases to larger than 1000 bases, but the specific concentration of RNA of various sizes is difficult to detect because the dissolved RNA concentration is too low, only at the picogram level. During the 6 experimental days, two relatively significant fluctuating periods in dissolved environmental RNA levels were observed (Figure 3c).

**Figure 3.**
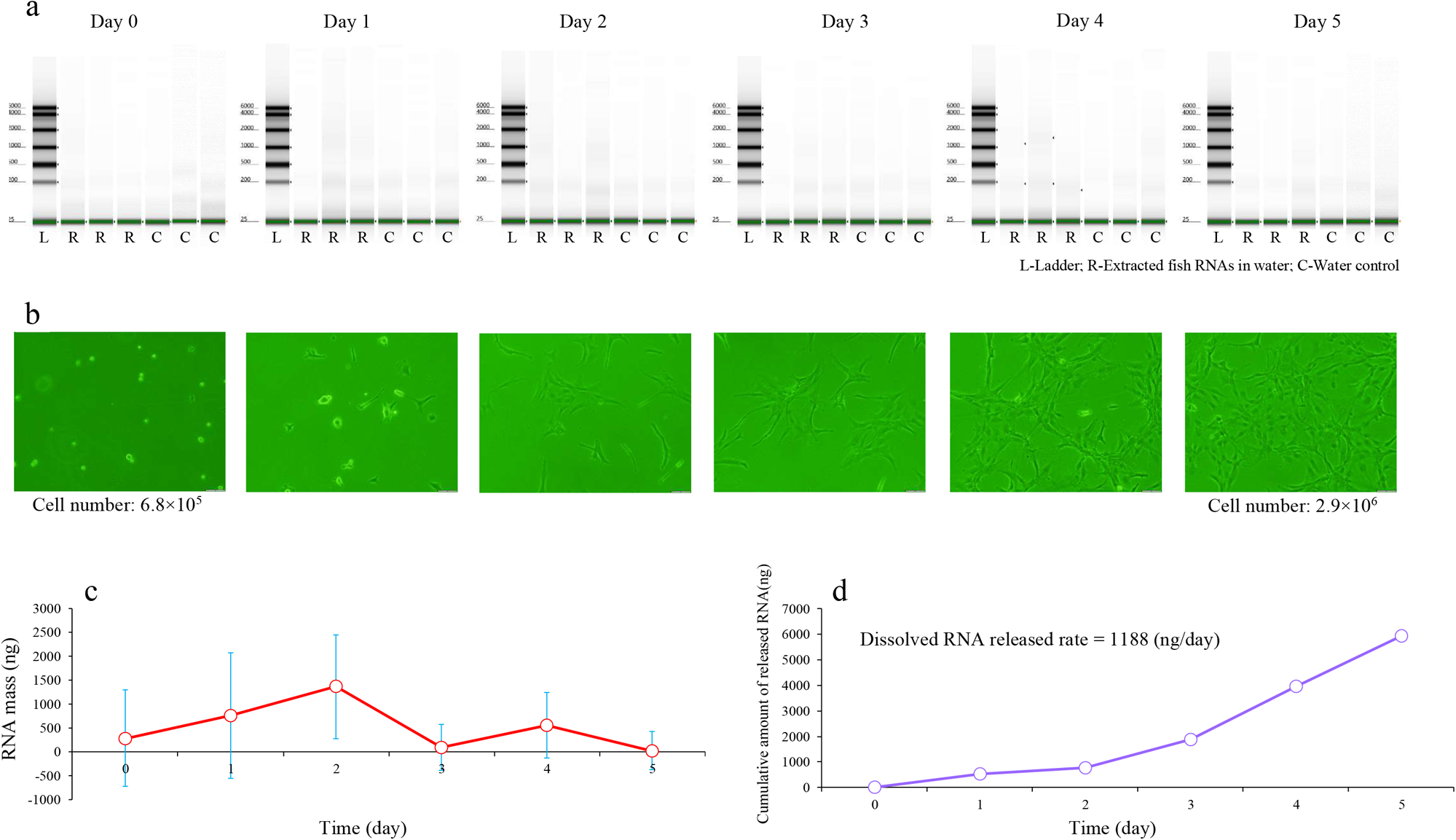
Dissolved RNAs released by living cells. In the figure, L represents the marker or the ladder, R represents extracted zebrafish RNAs in water, and C represents the water control. a. The results of detection by TapeStation. b. The physiological state of zebrafish BRF41 cells. c. The dissolved RNA mass in a flask. The solid pink curve represents the amount of dissolved RNA in the medium, and the blue bar is the standard deviation. d. The cumulative amount of dissolved RNAs released by living zebrafish BRF41 cells in a flask.

On Day 0, the media samples were collected 3 hours after the cells and media were placed in the flasks, and there were approximately 286 ng of dissolved environmental RNAs as a background level in a flask at the time of sampling (Figure 3c). According to the results calculated by the model to estimate gross transcription rate (Xu and Asakawa, 2023), without considering degradation, the calculated amounts of dissolved environmental RNAs released each day in an experiment flask showed fluctuations (Figure 3d): there were 0 ng, 525 ng, 262 ng, 1098 ng, 2084 ng, and 1973 ng on the 0^th^, 1^st^, 2^nd^, 3^rd^, 4^th^, and 5^th^ day, respectively.

The cells grew well, most cells attached to the bottom surfaces of the flasks, the cell shapes stretched on the 1^st^ day, and the number of cells increased by more than 3 times during the experiment (Figure 3b).

### 2.4 Dissolved environmental RNAs released by the cells in nuclease-free water

The rates and total amount of dissolved RNAs released from the cells in nuclease-free water could be detected (Figure 4). In total, each cell in nuclease-free water released approximately 4.2 pg RNA into the liquid environment during the 6-day experiment. During these 6 days, the dissolved environmental RNAs in water showed a fluctuating trend (Figure 4c).

**Figure 4.**
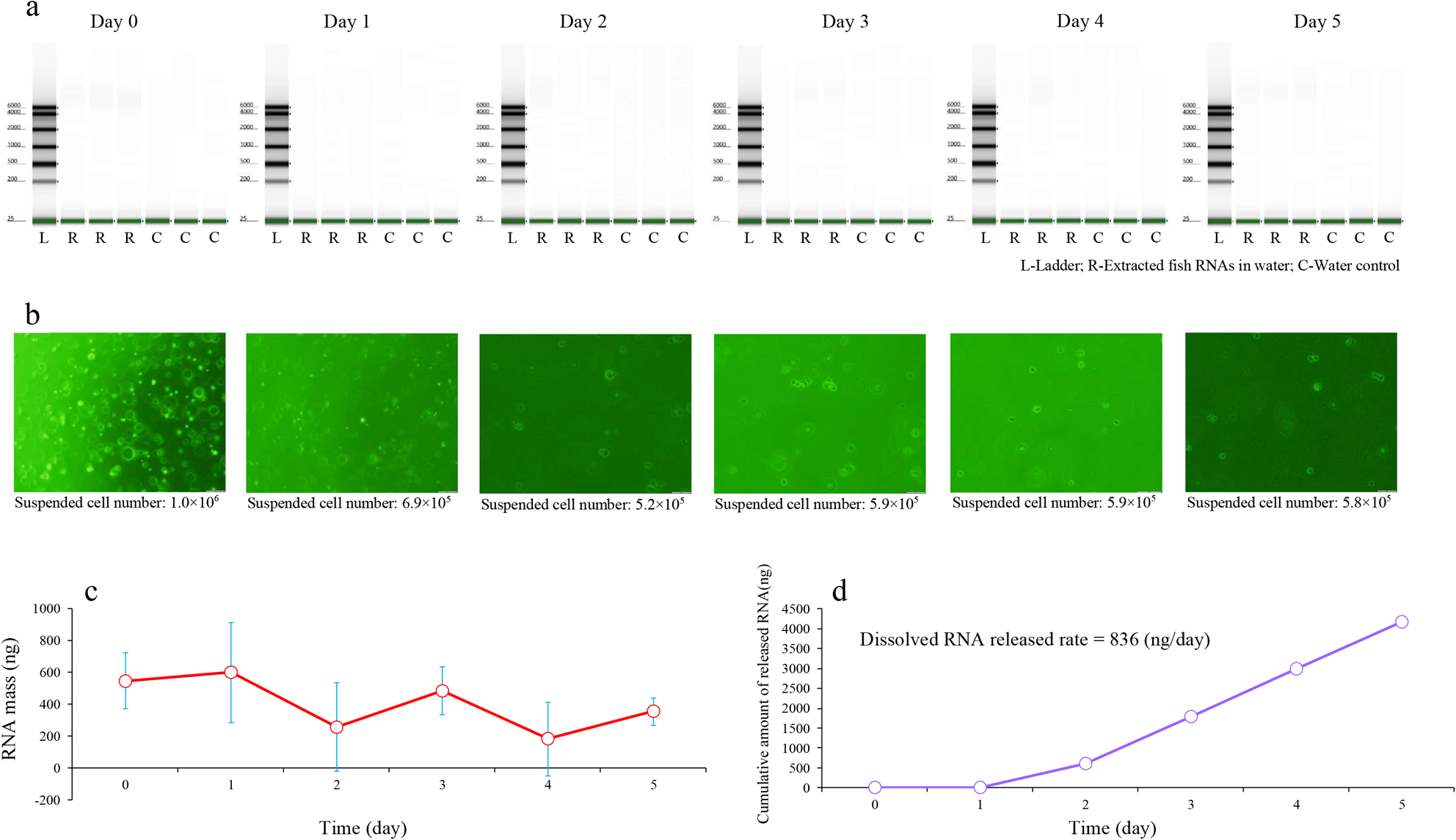
Dissolved RNAs released by the cells in nuclease-free water. In the figure, L represents the marker or the ladder, R represents extracted zebrafish RNAs in water, and C represents the water control. a. The results of detection by TapeStation. b. The physiological state of zebrafish BRF41 cells. c. The dissolved RNA mass in a flask. The solid pink curve represents the amount of dissolved RNAs in nuclease-free water, and the blue bar is the standard deviation. d. The cumulative amount of dissolved RNAs released by the cells in nuclease-free water in a flask.

On Day 0, the water samples were collected 3 hours after the cells and nuclease-free water were placed in the flasks, and there were approximately 546 ng dissolved environmental RNAs as a background level in a flask at the time of sampling (Figure 4c). According to the results calculated by the model to estimate the gross transcription rate (Xu and Asakawa, 2023), without considering degradation, the calculated amounts of dissolved environmental RNAs released each day in an experimental flask showed a rising trend (Figure 4d): there were 0 ng, 0 ng, 597 ng, 1195 ng, 1195 ng, and 1195 ng on the 0^th^, 1^st^, 2^nd^, 3^rd^, 4^th^, and 5^th^ days, respectively.

The cells were suspended in the water, and almost no cells were attached to the bottom surfaces and grew during the experiment (Figure 4b). Seventy percent of the suspended cells were observed in the flasks on the 1^st^ day, and 50-60% of the suspended cells could be observed on the 2^nd^ ∼ 5^th^ days. On the 3^rd^ day, cell fragmentation was observed.

### 2.5 Comparison of degradation rates of dissolved environmental RNAs under different conditions

The results of comparing the degradation rates of dissolved environmental RNAs showed that dissolved environmental RNAs in water from zebrafish cells degraded faster in the presence of zebrafish cells (Figure 5). Under the conditions without zebrafish cells, the survival rates per 10 days of zebrafish RNAs were relatively stable in the scope of 67∼92%/10 days, and the average survival rate per day was 98.4%/day (Figure 5a). Under living zebrafish cell conditions, the degradation of environmental zebrafish RNAs was low on the 1^st^ and 2^nd^ days, with survival rates per day of 100%/day and 93.8%/day, respectively, but degradation increased sharply after the 3^rd^ day, and the average survival rate per day on the 3^rd^, 4^th^, and 5^th^ days was 18.2%/day (Figure 5b). Under the conditions with dying zebrafish cells, the environmental zebrafish RNAs degraded gradually on the 1^st^ and 2^nd^ days, with survival rates per day of 80.7%/day and 32.5%/day, respectively; the degradation remained at relatively stable high levels after the 3^rd^ day, and the average survival rate per day of the 3^rd^, 4^th^, and 5^th^ days was 20.4%/day (Figure 5c).

**Figure 5.**
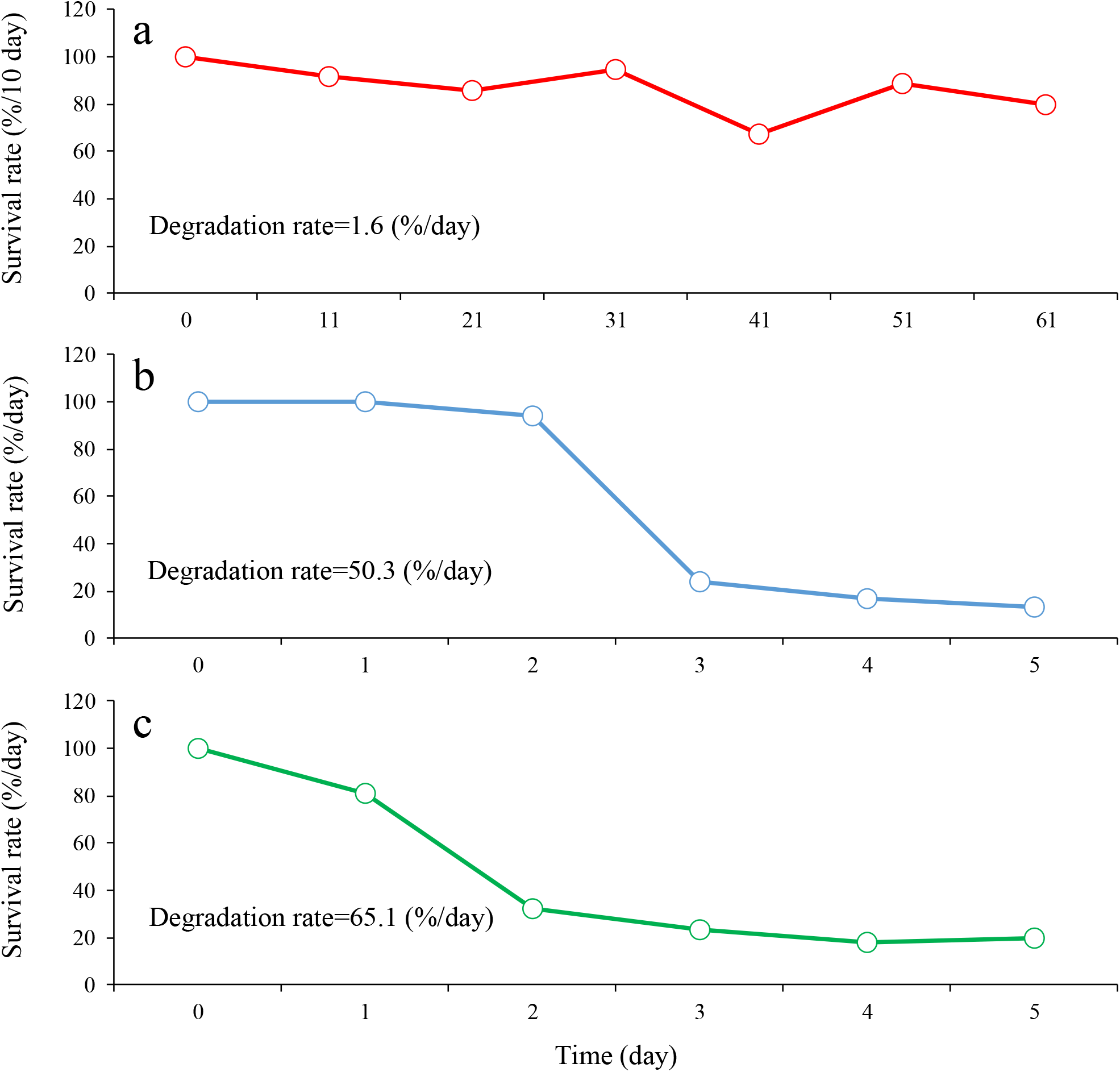
Comparison of degradation rates of dissolved environmental RNAs among different conditions. a. No cell: RNAs in water in the tubes. The red hollow circle is the survival rate per 10 days. The data were from the degradation of free RNAs in water in the tubes (Figure 2). b. Living cells: RNAs in flasks with zebrafish BRF41 cells in normal medium. The blue hollow circle is the survival rate per day. The data were from the experiment of dissolved environmental RNAs released by living cells (Figure 3). c. Dying cells: RNAs in flasks with zebrafish BRF41 cells in nuclease-free water. The green hollow circle is the survival rate per day. The data were from the experiment of dissolved environmental RNAs released by the cells in nuclease-free water (Figure 4).

### 2.6 Estimation of dissolved environmental RNA level fluctuations in a fish tank

In a study about environmental RNAs released by zebrafish (Jo et al., 2022b), the amount of a mitochondrial DNA gene CytB from DNA extraction from 300 ml water sample (from the tanks removing zebrafish) on the 0^th^ day was 1203.5∼8516.9 copies/PCR. Each adult zebrafish cell is reported to have 5500 copies of mitochondrial genomes (Artuso et al., 2012). Considering the degradation of cellular mitochondrial DNA in water, I hypothesized that each zebrafish in the water sample collected in the experiment (Jo et al., 2022b) had 1203.5∼5500 copies/PCR of the CytB gene, so each fish tank had 3∼116 cells/L released from zebrafish. The released zebrafish cells in the tanks with freshwater in that experiment were similar to the BRF41 zebrafish cells in nuclease-free water. Therefore, 12.6∼487.2 pg/L RNAs are released from zebrafish cells into the environment. The levels of dissolved environmental RNAs in water in each tank in that experiment fluctuated during the 5-day experimental period (Figure S1); the possible lowest level and possible highest level of dissolved environmental RNAs in water in each tank were 0.5 pg/L and 69.3 pg/L, respectively.

## 3. Discussion

### 3.1 The sizes of dissolved environmental RNAs in water

According to the present experiments (Figure 1, Figure 2, Figure 3, and Figure 4), RNAs with different sizes could exist in water as dissolved environmental RNAs. If the dissolved environmental RNAs in water included functional RNAs with different sizes, such as informative mRNAs, short noncoding RNAs, and long noncoding RNAs (Eddy, 2001; Fatica and Bozzoni, 2014), the transcriptomic data of related organisms were stored in the water samples. In fact, the RNA species and amounts of detectable dissolved environmental RNAs were limited: the dissolved environmental RNAs detected in the present study were mostly less than 500 nucleotides, which suggested that the detectable dissolved environmental RNAs in water were RNAs with small sizes and RNA degradation fragments. For dissolved environmental RNAs that may be widely present in natural water bodies, however, even data on trace amounts of RNA degradation fragments may provide important biological information. Further analysis of dissolved environmental RNAs will need sequencing and fragment assembly techniques, and related methods in ancient DNA seem to be helpful (Pääbo and Wilson, 1988; Pääbo, 1989; Willerslev and Cooper, 2005).

The length of RNA depended on the number of nucleotides. There is a rise per base pair of 0.25∼0.28 nm in the persistence length and an effective hydrodynamic diameter of approximately 3 nm of a helix RNA, and the flexibility of transient changes in molecular conformation may make the actual length of RNA longer or shorter (Arnott et al., 1967; Kebbekus et al., 1995; Hagerman, 1997). Until the current detection of environmental RNAs in water, the minimum pore size of filters for filtering water samples was larger than 0.2 µm (Wu and Liu, 2018; Wood et al., 2020; Hempel et al., 2022). In theory, naked RNAs with all lengths can pass through pore sizes larger than 0.2 µm in a certain conformation because they are only 3 nanometers in hydrodynamic diameter. Do RNAs with persistence lengths larger than the pore sizes of the filters, such as RNAs with more than 2000 nucleotides (0.50∼0.56 µm in the persistence length), sometimes fail to pass through the filters? A study showed that the large ribosomal subunit of *Alexandrium pacificum* could be partly collected in filters with pore sizes of 0.45∼5 µm (Zaiko et al., 2022). In the present study, however, there were no significant differences between the levels of zebrafish RNAs with different sizes, including the large ribosomal subunit, in the samples before and after filtration (P>0.05). This may mean that the persistence length of RNAs being larger than the pore size of the filter was not the reason why the filters could collect naked RNAs; other reasons need to be considered and verified. Some filter materials have the affinity of adsorption of RNAs, which might be one of the reasons (Sakano and Kamatani, 1992). It is also possible that during filtration, other particles first clog the pores of the filter, preventing RNA from passing through. RNA attached to other large particles would also fail to pass through the filter, but in this case, these RNAs are not dissolved environmental RNAs.

### 3.2 The release of dissolved environmental RNAs by living and dying cells

The results of the detectable dissolved RNAs in the environment of living zebrafish cells support the idea that living cells can release dissolved environmental RNAs, although it is also possible that dissolved environmental RNAs were released after some cells died. There have been some studies on the secretion of RNA by cells: RNA signals have been detected in the medium culturing mouse embryonic fibroblast 3T3 cells (Kolodny et al., 1972); RNAs can be secreted with extracellular vesicles as carriers, with the partial aim of cell‒cell communication (O’Brien et al., 2020; Veziroglu and Mias, 2020; Karimi et al., 2022); RNAs can also be secreted by binding to other macrobiomolecules (Sork et al., 2021). These studies support the results of the present research, but targeting dissolved RNAs secreted by living cells into aqueous environments is still lacking in previous studies. The present study measured the sustained secretion of dissolved RNAs from zebrafish cells and calculated the rate of dissolved RNA released by zebrafish cells by using the algorithms developed (Xu and Asakawa, 2023), and such work could help improve the techniques to directly detect dissolved RNAs released into liquid environments by living cells and quantitatively analyze the behaviors of dissolved environmental RNAs in water.

The dissolved environmental RNA abundance in the medium culturing living zebrafish cells fluctuated (Figure 3c). This fluctuation resulted from the dynamic relationship between the release of dissolved environmental RNAs by cells and degradation in the medium, such as some biomolecule fluctuations caused by the coupling of a positive factor and a negative factor (Elowitz and Leibler 2000). The fact that the RNA levels in cells under normal conditions usually fluctuate has been reported by some researchers (Deng et al. 2014; Briggs et al. 2018; Rodriguez et al. 2019; Xu and Asakawa, 2021). Did the fluctuations in the levels of dissolved environmental RNAs released by living zebrafish cells reflect the RNA level fluctuations in living zebrafish cells? This question cannot be answered with the current experimental data, especially without data on changes in cellular RNA levels over time. Cellular RNA level fluctuations in the related references (Deng et al. 2014; Briggs et al. 2018; Rodriguez et al. 2019; Xu and Asakawa, 2021) referred to RNA of specific genes, but the fluctuations in the release rate of dissolved environmental RNAs in the present study referred to total RNAs. Moreover, the design of sampling the medium per day in the present study might be a too long sampling time interval compared to the sampling design for the previous experiments of measuring cellular RNA levels.

Zebrafish cells cultured in nuclease-free water were regarded as dying cells because factors such as the osmotic pressure environment and nutrient supply make it difficult for zebrafish cells to survive. The dissolved environmental RNA abundance in the flasks containing dying cells fluctuated (Figure 4c). The standard deviation of the dissolved environmental RNA abundance in the flasks containing dying cells during the experimental period and that of living cells was 116.2 and 500.6, respectively, showing that the dissolved environmental RNA abundance in the flasks containing dying cells had lower fluctuations. This may indicate that the degree of fluctuations of dissolved environmental RNA in water can provide useful information to identify living and dead cells in water. The daily released amounts of dissolved environmental RNAs by the dying cells showed two plateaus from 0 ng/flask to 1195 ng/flask. The former low plateau period may be that under high environmental pressures, the function of the cell membrane to release RNA outside the cell was temporarily blocked; in the latter high plateau period, the permeability of the cell membrane increased, and the cell leaked RNAs at a steady rate. In necrosis, one kind of cell death, when a cell under environmental damage presses, its cell membrane might lose permeability at first, and after some cell biological steps, the cell becomes leaky, releasing its content into the surrounding environment (D’Arcy, 2019). It seemed that the process of necrosis was similar to the results of RNAs released by zebrafish cells in nuclease-free water. Whether nuclease-free zebrafish cells undergo necrosis is worth further study.

Using the present experimental results of dissolved environmental RNAs in water released from zebrafish cells, the dissolved environmental RNA level fluctuations of 0.5∼69.3 pg/L in a fish tank reported in a study (Jo et al., 2022b) could be calculated (Figure S1), and such low concentrations of RNAs in water could not be detected by using the current techniques. Although this calculation result of dissolved environmental RNAs in water might be different from real water bodies due to the lack of detailed impacts of the complex physical, chemical, and biological factors in the field, the estimation could provide some information on cases that could not be obtained in real work under the current techniques.

The limitation of method protocols in the present study may affect the results of dissolved environmental RNAs released by cells. The quantitative range of High sensitivity RNA ScreenTape^@^ for measuring dissolved environmental RNAs was 500∼10000 pg/μL, but there were some detection results in the present experiments out of this range, which might lead to the results of fluctuated levels of RNA. In the experiment of degradation of free RNAs in water in the tubes, the degradation rate of RNAs in one repeat was much lower than the other two, and contamination by exogenous RNAs may not be the reason because there was almost no RNA contamination in the three control groups (Figure 2c); thus, the real reason for this result was unknown. Moreover, using the algorithm (Xu and Asakawa, 2023) to estimate the total amount and the release rate of dissolved environmental RNAs released by the cells requires a prerequisite: RNAs are released independently of the levels of dissolved environmental RNAs in liquid environments, whereas the degradation rate of dissolved environmental RNAs is proportional to the levels of dissolved environmental RNAs. There is no detailed evidence to explain whether the results of the present study are strictly consistent with this prerequisite. Therefore, improved protocols are urgently needed.

### 3.3 The degradation of dissolved environmental RNAs outside the cells in water

The half-life of RNA in cells is usually a few minutes to 2 hours in nonmammal cells and a few minutes to 48 hours in mammalian cells (Petersen et al., 1976; Cai and Winkler, 1993; Deng et al., 2013; McManus et al., 2015; Baudrimont et al., 2017; Steiner et al., 2019; Yamada and Akimitsu, 2019). RNAs that survive too long and accumulate in excess are harmful to cells; therefore, after an RNA is transcribed, its lifespan is controlled by the RNA surveillance system and degradation system (Cole et al., 1992; Isken and Maquat, 2007; Houseley and Tollervey, 2009). The cellular factors for surveilling and degrading RNAs in cells include endoribonucleases, exoribonuclease, the TRAMP complex, upframeshift proteins, microRNAs, etc. (Frischmeyer et al., 2002; Behm-Ansmant and Izaurralde, 2006; Chang et al., 2007; Deana et al., 2008; Eulalio et al., 2009; Houseley and Tollervey, 2009; Sohrabi-Jahromi et al., 2019). These factors work precisely and efficiently in cells to degrade RNA in time to ensure cell health. When RNAs are released outside the cells to be dissolved environmental RNAs in water, the cellular systems to surveille and degrade RNAs cannot work. The factors that induce RNA degradation outside cells are different from those in cells. RNAs outside the cells are thought to be readily degraded by ribonuclease (RNase) released into the environment by various organisms (Harder and Schröder, 2002; Rosenberg 2008; Rossier et al., 2009; Luhtala and Parker, 2010). Other biological, physical, and chemical factors, such as hydrolytic enzymes, aqueous solutions, and ions, also play an important role in the degradation of RNAs outside the cells (Tenhunen, 1989; Matsumura and Komiyama, 1997; AbouHaidar and Ivanov, 1999; Chatterjee et al., 2022; Zhang et al., 2023). However, factors that degrade RNA outside the cell appear to have lower RNA degradation efficiency than the RNA surveillance and degradation system inside the cell. The half-life of particle environmental RNAs from nonmammalian organisms in water, which are collected in filters, is reported to range from 8∼68 hours (Wood et al., 2020; Marshall et al., 2021; Jo et al., 2022b; Kagzi et al., 2022), longer than the half-life of RNA inside nonmammalian cells and longer than that in most mammalian cells.

The experiment of the degradation process of free RNAs in water in the present study was carried out in a nonsterile environment, but the trace RNA level in the controls indicated that there was almost no RNA contamination from water, air, and tubes, so the degradation process of free RNAs in water under a relatively open environment could be studied in this experiment (Figure 2). Some RNAs in the groups of RNAs in water could still be detected after 2 months in the tubes, and the half-lives of RNAs were 20∼43 days, much longer than the half-lives of RNAs inside cells and half-lives of particle environmental RNAs described above. On the one hand, the water in the tubes might contain fewer factors that degraded RNAs compared to the environments inside cells and in open water bodies, leading to low RNA degradation rates. On the other hand, the definition of RNA degradation used in the present study was slightly different. In some studies (McManus et al., 2015; Baudrimont et al., 2017), whether the RNA has been degraded is judged by whether a certain length (especially a whole length) of RNA sequence is detected; that is, RNA degradation is an all-or-none phenomenon. However, this definition might be insufficient for practical applications. For example, an RNA missing one base sometimes has the same function as the original RNA, or only the edited RNA has an important function. If RNA degradation is defined as the judgment of whether a base is losing, then the current experimental results were difficult to estimate the half-life of RNA. Instead, the present study regarded RNA degradation as a process in which the total mass of RNA decreases; thus, the RNA half-life was the time it took for the total mass of RNA to be reduced by half. Smaller and smaller RNA degradation fragments were observed as the experiment time was prolonged, as shown by different detection methods (Figure 2). The type of RNA in the tubes and the degradation properties of the individual RNAs could also affect the estimation of overall RNA half-lives. The quantitative relationship between RNA fragment length, experimental time, and environmental factors was still difficult to estimate in the current experimental data, which requires further research.

Dissolved environmental RNAs from zebrafish degraded faster in water with zebrafish cells than in water without zebrafish cells (Figure 5). High degradation rates of dissolved environmental RNAs in the water environments with living cells indicated that the living cells might secrete some factors into the medium to degrade RNAs released by themselves; according to the shape of the survival rate curve, these degradation factors were initially low but accumulated in increasing amounts (Figure 5b). In the case of water environments with dying cells, the related factors released by the broken cells might play an important role in the degradation of dissolved environmental RNAs, resulting in higher degradation rates (Figure 5c). In the experiment of free RNAs in water in the tubes (Figure 5a), the tubes had enzymes secreted by the bacteria to breakdown the RNA, but how the activity of these enzymes on zebrafish RNA degradation compared to the activity of related enzymes in zebrafish cells was unknown. RNA can be adsorbed on the surface of cells and small particles (Sakano and Kamatani, 1992; Yu et al., 2013; Lardeux et al., 2022), which will also reduce the amount of dissolved environmental RNAs in water with zebrafish cells. The RNAs mentioned in Figure 5a, Figure 5b, and Figure 5c were intracellular RNAs, the RNAs secreted by living cells, and the RNAs secreted by cells and released by cell rupture, respectively, and the difference in the source of RNA also caused the difference in RNA degradation rates. Further research on the degradation of different RNA sequences and the degradation specificity of different RNA degradation factors will provide a reference for answering related questions.

## 4 Conclusion

The present study showed that dissolved environmental RNAs could come from living cells and dying cells. The amount of dissolved environmental RNAs released from living zebrafish cells per day accounted for approximately 1∼7% of the cellular RNA mass. Further analysis of these dissolved environmental RNAs with different sizes can provide useful information for research such as noninvasive health detection of aquatic organisms and RNA communication between cells and/or individuals in water. The dying zebrafish cells released dissolved environmental RNAs at different rates from those of living zebrafish cells.

The low degradation rates of dissolved environmental RNAs allow them to remain in water for a longer time than previously estimated. RNA is more easily degraded inside cells, including in the liquid environment in which the same cells exist, than in water. The dissolved environmental RNAs in the water could be detected for more than two months at room temperature, although the most detectable dissolved environmental RNAs might be degraded as small fragments of RNA sequence. Thus, dissolved environmental RNA may be one of the trace background chemical components in some water bodies, and it can retain the information of the organisms that once existed or currently exist for a certain long period of time.

However, attention should be given to the problems of detection protocols in the present study. For example, some detection results of dissolved environmental RNAs in the present study were out of the quantitative range of High sensitivity RNA ScreenTape^@^ for measuring dissolved environmental RNAs; in the experiment of degradation of free RNAs in water in the tubes, the standard deviation values of the results detected by NanoDrop were too high (Figure 2c). Thus, improved protocols for the detection of dissolved environmental RNAs in water in the field are urgently needed. Despite these problems, the present study may provide information to develop new ways to measure the dynamics of dissolved environmental RNAs in water and quantitatively analyze RNAs released to liquid environments by living and dying cells.

## 5. Methods and materials

### 5.1 Culture of zebrafish BRF41 cells and RNA extraction

Zebrafish BRF41 cells (Cell number: RCB0804) were purchased from Cell Bank, RIKEN BioResource Research Center, Japan, and cultured in 25 mL or 75 mL flasks with 84% Leibovitz’s L-15 medium, 15% fetal bovine serum, and 1% 10 mM HEPES–NaOH (Tamura et al., 2006). The culture temperature was 33 ℃. The frequency of cell subculture was once per 10 days, with trypsin to suspend cells from flask bottoms and phosphate buffered saline to wash cells. RNA in zebrafish BRF41 cells was extracted with TRIzol^TM^ reagent.

### 5.2 Free RNAs through the 0.45 µm filter

RNA extracted from zebrafish BRF41 cells was diluted with nuclease-free water to a concentration of 65 ng/µL. There were three repeats of RNA solution and three controls of nuclease-free water. The liquid was passed through a 0.45 µm filter unit (Sterivex^TM^, EMD Millipore Corporation, Germary), and the filtrate was collected. The concentration and fragment sizes of the RNAs in the liquid before and after filtration were measured by a NanoDrop ND-1000 Spectrophotometer (Marshall Scientific LLC) and RNA ScreenTape system (Agilent 2200 TapeStation, Agilent Technologies, Inc.).

### 5.3 Degradation of free RNAs in water in the tubes

RNA extracted from zebrafish BRF41 cells was diluted with tap water to a concentration of 65 ng/µL. The liquid was placed into 1.5 mL centrifuge tubes at room temperature (25 °C). The centrifuge tubes were capped unless they were being used for sampling. There were three repeats of RNA solution and three controls of tap water. The RNA concentrations were measured by a NanoDrop ND-1000 spectrophotometer, normal gel electrophoresis (2.5% agarose), and RNA ScreenTape system. To simulate the water environment in which microorganisms, dust, etc., may enter, the tubes were placed in a nonsterile environment, and the sampling operation process was carried out in a nonsterile environment.

### 5.4 Dissolved environmental RNAs released by living cells

Zebrafish BRF41 cells were cultured in 15 ml L-15 medium in 75 mL flasks. The number of seed cells was 6.8×10^5^. There were three repeats of zebrafish BRF41 cells cultured in 15 ml L-15 medium (called “Cells in Medium”) and three controls of L-15 medium without zebrafish BRF41 cells (called “Medium Control”). After subculture, the RNAs in the medium from the Cells in Medium and Medium Control groups were measured on the 0^th^ day, 1^st^ day, 2^nd^ day, 3^rd^ day, 4^th^ day, and 5^th^ day. One milliliter of medium was removed and passed through a 0.45 µm filter unit, and the filtrates were collected. The concentration and fragment sizes of the RNAs in the filtrates were measured by High Sensitivity RNA ScreenTape^@^ (Agilent 2200 TapeStation, Agilent Technologies, Inc.) and NanoDrop ND-1000 Spectrophotometer. The cells in the flasks were observed under a microscope (Olympus IX70) and photographed using the CellSens standard. On the 5^th^ day, the cell numbers in the cells in medium flasks were counted under the normal subculture procedure. The experimental operation was carried out in a sterile environment.

The RNA mass in a flask of cells in medium was calculated as follows:

RNA mass in a flask = (RNA concentration in Cells in Medium - RNA concentration in Medium Control) × volume of medium in the flask

The RNA mass was not the total amount of dissolved RNAs released by the living cells during the experimental period but only a snapshot of RNA abundance. The model to estimate gross transcription rates from RNA level fluctuation data (Xu and Asakawa, 2023) was used to estimate the total amount and release rate of dissolved RNAs released by living cells.

### 5.5 Dissolved environmental RNAs released by the cells in nuclease-free water

Zebrafish BRF41 cells cultured in L-15 medium were suspended from flask bottoms by using trypsin, rinsed with phosphate buffered saline, resuspended in L-15 medium, and then placed into flasks filled with 15 ml nuclease-free water. The number of seed cells was 1.0×10^6^. There were three repeats of zebrafish BRF41 cells cultured in nuclease-free water (called “Cells in NF”) and three controls of nuclease-free water without zebrafish BRF41 cells (called “NF Control”). The RNAs in the water from Cells in the NF and NF control groups were measured on the 0^th^ day, 1^st^ day, 2^nd^ day, 3^rd^ day, 4^th^ day, and 5^th^ day. One milliliter of nuclease-free water was removed and passed through a 0.45 µm filter unit, and the filtrates were collected. The concentration and fragment sizes of the RNAs in the filtrates were measured by High Sensitivity RNA ScreenTape^@^ and NanoDrop ND-1000 Spectrophotometer. The cells in the flasks were observed under a microscope (Olympus IX70) and photographed using the CellSens standard. Every day during the experiment, the numbers of suspended cells in the cells in medium flasks were counted with a hemocytometer. The experimental operation was carried out in a sterile environment.

The RNA mass in a flask of Cells in NF was calculated as follows:

RNA mass in a flask = (RNA concentration in Cells in NF - RNA concentration in NF Control) × volume of nuclease-free water in the flask

The model to estimate gross transcription rates from RNA level fluctuation data (Xu and Asakawa, 2023) was used to estimate the total amount and release rate of dissolved RNAs released by the cells in nuclease-free water.

### 5.6 Estimation of dissolved environmental RNA level fluctuations in a fish tank

The level of dissolved environmental RNA in a fish tank was difficult to directly detect. Cells in nuclease-free water were dying cells, and they were similar to the cells released by aquatic organisms in water. The experimental data of a study on environmental RNAs in water contributed by zebrafish (Jo et al., 2022b) were selected, combined with the results of dissolved environmental RNAs released by the cells in nuclease-free water in the present study, to calculate dissolved environmental RNA level fluctuations in a fish tank.

## Acknowledgments

This work was supported by JST SPRING, Grant Number JPMJSP2108.

## Supplementary figures

**Figure S1.**
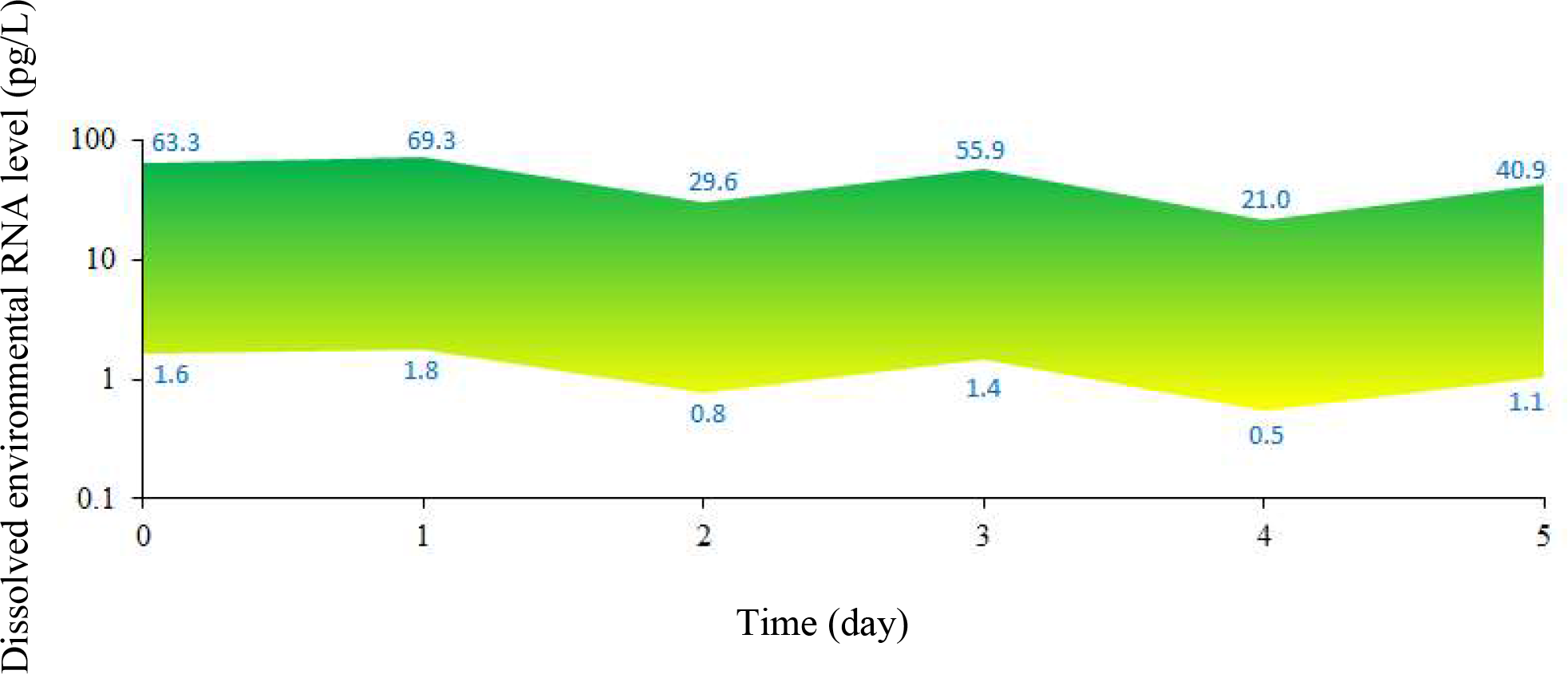
The range of the dissolved environmental RNA levels in a fish tank removing zebrafish. Yellow represents lower dissolved environmental RNA levels, and green represents higher dissolved environmental RNA levels. The data of environmental DNA of the mitochondrial genome and mitochondrial genome copies of zebrafish are from the references (Artuso et al., 2012; Jo et al., 2022b). After the calculation of the data, each fish tank had 3∼116 cells/L released from zebrafish. The data of dissolved environmental RNA amount released by zebrafish cells were from the experiment of dissolved environmental RNAs released by the cells in nuclease-free water (Figure 4).

